# Using VR and eye-tracking to study attention to and retention of AI-generated ads in outdoor advertising environments

**DOI:** 10.1101/2024.08.15.607684

**Authors:** Sue Lim, Hee Jung Cho, Moonsun Jeon, Xiaoran Cui, Ralf Schmälzle

## Abstract

In contemporary urban environments, advertisements are ubiquitous, capturing the attention of individuals navigating through cityscapes. This study simulates this situation by using VR as an advertising research tool and combining it with eye-tracking to rigorously assess attention to and retention of visual advertisements. Specifically, participants drove through a virtual city with 40 AI-generated, experimentally manipulated, and randomly assigned advertisements (20 commercial, 20 health) distributed throughout. Our results confirm theoretical predictions about the link between exposure, visual attention, and incidental memory. Specifically, we found that attended ads are likely to be recalled and recognized, and concurrent task demands (counting sales signs) decreased visual attention and subsequent recall and recognition of the ads. Finally, we identify the influence of ad placement in the city as a very important but previously hard-to-study factor influencing advertising effects. This paradigm offers great flexibility for biometric advertising research and can be adapted to varying contexts, including highways, airports, and subway stations, and theoretically manipulate other variables. Moreover, considering the metaverse as a next-generation advertising environment, our work demonstrates how causal mechanisms can be identified in ways that are of equally high interest to both theoretical as well as applied advertising research.

## 1. Introduction

Picture yourself navigating through the bustling streets of a city. As you pass by storefronts and tall office buildings, you see signs and letterboxes, trees, and streetlights. Occasionally, you also look at colorful advertisements, like the one by an intersection that features hip furniture and says: “Don’t miss the biggest sale of the season!” Another billboard, placed by a small park, is a PSA about smoking cessation. Two basic questions are: 1. What determines which ads draw your attention? 2. What will you remember at the end of your drive?

This scenario describes a typical situation in real-world advertising. The two questions are about as old as advertising research itself (Kreshel, 1993), and the theoretical concepts of attention and retention (memory) have been part of advertising for over a century (Hopkins, 1923; Karslake, 1940; Nixon, 1924; Poffenberger, 1925). Virtually all theories of advertising, information processing models from psychology, and insights from neuroscience support the idea that attention to an ad is an important first step toward its effectiveness (Lang, 2009; Lindsay, 2020; Rodgers and Thorson, 2012; Rossiter and Bellman, 2005), and numerous empirical studies have measured the predictors and consequences of attention to ads, which includes ad retention.

Broadly speaking, attention is thought to be determined by (i) characteristics of the ad itself (e.g., attractive images), (ii) the context in which it is displayed (e.g., among other ads vs. alone), or (iii) receiver factors (e.g., distraction, motivation). Once an ad has attracted attention, it is more likely to leave a trace in the observer’s memory (McGuire, 2001; Sherman and Turk-Browne, 2022). The practice of ad pretesting further underscores the role of attention and memory as key advertising factors: Leading agencies rely on metrics of attention (from self-report to biometrics like eye-tracking) and memory (most notably recall and recognition; Krishnan and Chakravarti (1999); Ostlund and Clancy (1982); Smit and Neijens (2011); Wedel and Pieters (2017)).

However, there remains a sizeable gap between the everyday city-driving scenario described above and the current practice of advertising research on attention and memory. In the real world, people can freely explore their environment and decide in a self-directed manner which ads and other objects in the environment to pay attention to. In addition, memory typically forms incidentally (Castel et al., 2015). By contrast, many advertising researchers control which ads participants see and artificially reduce the environment, sometimes even down to display only a single ad in isolation. Such constraints are driven by the need to experimentally manipulate isolated theoretical variables, control confounds, and standardize conditions. Advertising textbooks and research methods courses discuss the underlying issues with these research paradigms at length. Yet, the gap between theoretical advertising research and practice remains one of the key challenges for the field. An ideal solution involves the integration of experimentation and measurement with the realism of field studies (Fitts and Hewett, 1977; King and Tinkham, 1989; Wilson et al., 2015) and experimental control while enhancing cost-effectiveness, scalability, and speed.

The main contribution of this paper is to showcase a novel approach that offers a promising solution to close this gap. Specifically, we leverage virtual reality (VR) with eye-tracking to study advertising effects in a realistic urban outdoor context. In doing so, we manipulate theorized factors under controlled settings, capture biometric indices of attention in real-time, and examine their effects on subsequent memory (see Figure 1 for the overview of our research paradigm).

**Figure 1:**
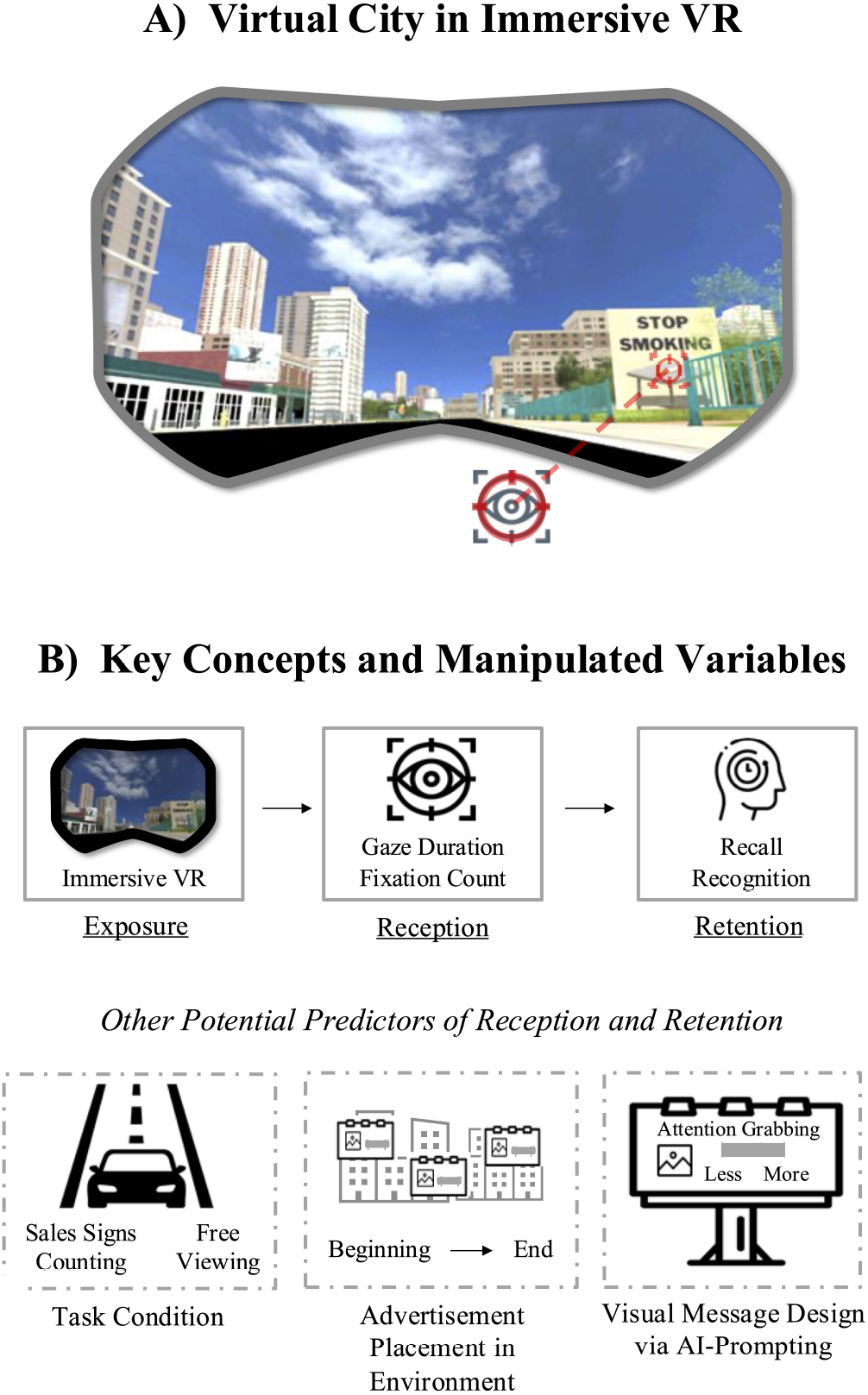
Overview of the VR City-Driving Paradigm. Participants were immersed in a virtual city via a VR headset integrated with eye-tracking capabilities (A). Our research paradigm examined the link between exposure, attention, and memory, as well as other predictors (B)

In this paper, we first introduce the theoretical concepts of attention and memory and the existing operationalization of the concepts. We then discuss the limitations of existing advertising effects research paradigms that use biometric measures and argue that there is a need for research that can measure exposure to, attention towards, and retention of advertisements in realistic contexts. From this, we conclude that the combination of VR and eye-tracking offers a powerful and flexible way to pre-test ad candidates in realistic environments. Next, we introduce the current study along with its hypotheses and research questions. Finally, we present the results and discuss their implications for advertising research and practice.

## 2. Theoretical Background: Attention and Memory in Advertising Research

Attention is a central concept in advertising, both for theoretical mechanisms and market impact (Davenport and Beck, 2001; Nixon, 1924). Although the term attention has many meanings that vary slightly across contexts and research areas, this paper focuses on selective attention to aspects of the external world - specifically messages presented in a city amidst other city-typical objects (Chun et al., 2011). This type of attention is referred to as overt visual attention, or the explicit deployment of eye gaze (via fixation) to select a subset of one’s visual surroundings for prioritized processing, which is typically associated with at least minimal conscious awareness (Lindsay, 2020).

In terms of measures of attention to ads, a variety of options exist - from retrospective self-report to in-the-moment think-aloud protocols to physiological measures like EEG or eye-tracking. Eye-tracking is the most valid and promising measure for this study, as it examines overt visual attention (Wedel and Pieters, 2008). Eye-tracking can answer questions like the following: Do people look at all at a specific message? Where, when, and for how long do they look? Does overt attention predict memory? These questions have considerable theoretical and practical significance for advertising research. Indeed, eye-tracking has already become the de-facto standard for the biometric quantification of attention in advertising (Fox et al., 1998; Huddleston et al., 2015; Wedel and Pieters, 2017), especially since the equipment has become commodified and easy to use.

Memory plays a crucial role in translating visual attention to an advertisement into tangible outcomes. While advertising objectives often extend beyond mere recall and aim to influence attitudes or drive purchase behaviors, an advertisement must imprint itself in the audience’s memory for it to have an effect. Accordingly, measures of ad memory, typically recall or recognition, have long been part of advertising research (Krishnan and Chakravarti, 1999; Smit and Neijens, 2011). Recall involves the active retrieval of a memory trace, whereas recognition only requires classifying an ad as having been seen (Loftus and Loftus, 2019).

Previous research in cognitive psychology reveals a link between attention and memory (Craik, 2002; Sherman and Turk-Browne, 2022; Uncapher et al., 2011). This attention-retention link is very compatible with advertising insights and motivates advertisers to achieve optimal exposure and wide reach, increasing the likelihood of advertisement success (McGuire, 2001; Rossiter and Bellman, 2005). Both attention allocation as well as memory formation are usually incidental: People look at ads in a self-directed manner, and they remember them as a by-product of their attention to the ad (Breuer and Rumpf, 2012; d’Ydewalle et al., 1988; Shapiro and Krishnan, 2001). Thus, understanding the connection between attention and incidental memory in advertising research is pivotal for enhancing message design strategies and effectively engaging customers. Although conducting such research in a laboratory setting is embedded with challenges, the virtual reality (VR) city paradigm introduced in this study offers a novel springboard that aids in overcoming limitations and simulating more realistic consumer environments.

### 2.1. VR and Eye-Tracking to Examine Attention and Memory in Simulated Environments

Advertising research has thoroughly examined the link between attention and retention of ads post-exposure, but methodological challenges persist. In particular, researchers seeking to pre-test ads or examine theoretical predictions in realistic environments face a significant challenge. Laboratory studies using stationary eye-trackers or screen-based paradigms (Boerman et al., 2015; Fischer et al., 1989; Lee and Ahn, 2012) offer experimental control, but they lack ecological validity. This lack of ecological validity limits their applicability to real-world contexts (De Pelsmacker, 2021; Vargas et al., 2017). While digital advertising has made strides in optimization through A/B testing (Kohavi and Thomke, 2017), such methods are unsuitable for research about exposure in physical locations such as a city. Conversely, field studies, with mobile eye-tracking devices or cameras provide realism but are costly, time-consuming, and subject to many confounds (AlKheder, 2024; Wilson and Casper, 2016; Xie et al., 2024).

An ideal research paradigm would allow researchers to simulate realistic message exposure conditions and environments but with exquisite experimental control and manipulation potential. Such conditions involve presenting the ads themselves, as well as contextual information such as where and when they appear and under which state viewers encounter them (e.g., distracted or fully engaged with the environment). Moreover, combining realistic viewing conditions with objective eye-tracking measures would be desirable, given the well-known advantages of biometric measures over self-report (Beard et al., 2024; Potter and Bolls, 2012; Read et al., 2018).

Therefore, we propose a solution that leverages VR with integrated eye-tracking capabilities. VR allows researchers to simulate any advertising context without being restrained by physical or financial limitations. For instance, researchers can create a virtual city and have people drive down its streets without having to close down a street, set up and pay for test billboards, or worry about regulations. Importantly, VR conveys a real sense of visuospatial presence and prompts immersed users to exhibit natural visual and attentional exploration behavior. With eye-tracking capabilities integrated into VR headsets, researchers can then gather real-time biometric data about people’s engagement with ads (Meißner et al., 2019; Schmälzle et al., 2023). Through this integration of eye tracking and VR, we can thus delve into the cause-effect sequence of information transfer from the ad in the environment into the audience’s visual system and subsequently to their storage in memory.

Similar reasoning has already motivated some prior work on advertising effects in simulated environments. For instance, Clay et al. (2019) examined navigation and gaze behavior in a virtual city that users can navigate freely. Kang et al. (2023) conducted a study in a virtual mall and found that attention to ads is associated with subsequent effects, and Wang et al. (2019) describes a system and pilot test for similar use cases. Bonneterre et al. (2024), another relevant recent study, built realistic urban settings to examine the user reactions to health messages. Similarly, Schmälzle et al. (2023) and Cho et al. (2024) built realistic highways to examine incidental exposure and attention. Beyond these directly related works, other research exists in related domains, such as VR-based eye-tracking to study shoppers’ attention in virtual supermarkets, tourists in virtual museums, or a variety of basic science questions in psychology and neuroscience (Anderson et al., 2023; Bischof et al., 2023; Moreno-Arjonilla et al., 2024).

## 3. The Current Study

Our study aims to rigorously examine the factors influencing attention and retention in a realistic outdoor advertising context by leveraging immersive VR integrated with eye-tracking and generative artificial intelligence (AI). Specifically, we simulated a common exposure environment (i.e., city) by creating a virtual city with 40 advertisements placed throughout. We varied the visual features of the advertisement with the help of generative AI to create two versions of the 40 advertisements (less vs. more attention-grabbing). Bringing all the features together, we examined how the advertisements are placed in the virtual environment, the assigned task of the participants, and the visual message design features (determined with the help of generative AI) influenced advertisement reception and retention (see Figure 2).

**Figure 2:**
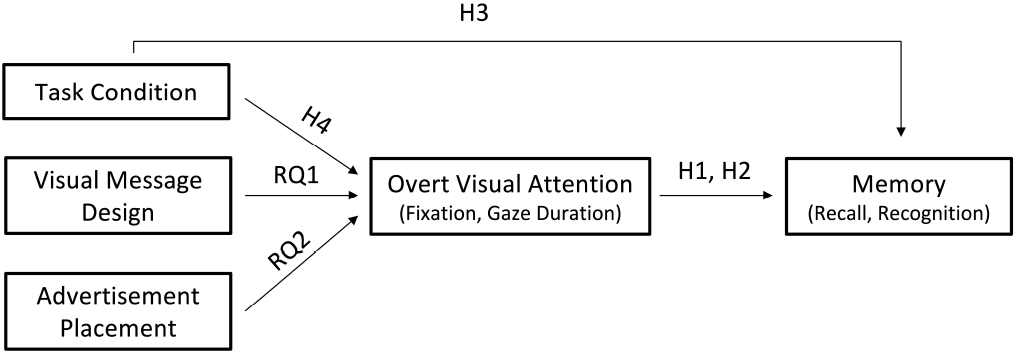
Conceptual Overview of the Predicted Links Between Attention to and Retention of Ads in Outdoor Contexts.

As discussed above, attention is a prerequisite for subsequent memory. Simply put, ads that are not looked at have no chance to make it into memory. Eye-tracking provides a sensitive, valid, and reliable measure of overt attention to ads (Bott et al., 2017; Bucher and Schumacher, 2006; Chang and Choi, 2014), and memory can be measured via recall and recognition tests (Postman et al., 1975). Among various measures of gaze behaviors, the current study delves specifically into fixations level (fixated 0 times, median count, or more than median count) and gaze duration (the total length of time that gaze persists post-fixation on a given stimulus). Fixation refers to gazing at the ad for a minimal, stable period. Prior eye-tracking data regarding visual information indicate both a positive correlation and a causal relationship between attention measures such as fixation level and subsequent recall/recognition tasks (Fehlmann et al., 2020; Olsen et al., 2016). More recent empirical findings on consumer behavior research using the eye-tracking method reveal that product exposure captured by fixation duration was strongly associated with brand recognition and memory performance (Kongmanon and Petison, 2022; Ronft et al., 2023).

Based on the reasoning laid out above, we posit

*H1: Fixation level will increase the likelihood of advertisement recall (H1a) and recognition (H1b)*.

*H2: Gaze duration will increase the likelihood of advertisement recall (H2a) and recognition (H2b)*.

Next, it is well known that attention faces critical bottlenecks and capacity limitations (Lang, 2009; Marois and Ivanoff, 2005). For example, focusing intensely on achieving a goal (e.g., counting certain signs) takes people’s attention away from accomplishing other tasks (e.g., remembering the content of non-target ads). In fact, literature on completing parallel tasks provides further evidence of a significant decline in performance when individuals attempt to handle multiple tasks simultaneously. Studies have shown that dual-task interference leads to a substantial reduction in recall (Armstrong and Chung, 2000; Voorveld, 2011) and recognition (Zhang et al., 2010), highlighting the competition for processing resources and the impact of divided attention. This illustrates the difficulties in managing parallel tasks and the significant cognitive load imposed on individuals, which in turn affects the overall performance. We also deduce that those occupied with a parallel task meant to distract them from the ads will spend less time looking at the ads (Kircher and Ahlstrom, 2017; Wolfe, 1998; Wolfe et al., 2022).

*H3: Engaging in a parallel task will decrease the likelihood of recall (H3a; recognition - H3b)*.

*H4: Engaging in a parallel task will decrease fixation (H4a) and gaze duration (H4b) on advertisements*.

In addition to the hypotheses, we pose two research questions about potential predictors of overt attention. We first tested the effects of visual features of the ad (more vs. less attention-grabbing) on overt attention (fixation level and gaze duration). Since many ad features could influence attention and researchers cannot experimentally control all confounding factors, we conducted prompt engineering with AI text-to-image generators to set a standardized conceptualization of more vs. less attention-grabbing ad images. We then used research-based principles about visual attention (Cho and Suh, 2020; Hammond, 2011; Liu et al., 2021) to select the appropriate generated image and integrate it with the background and ad text (see the Methods section for more details). Some existing studies have observed people’s overt attention to certain aspects of the ads or messages (Lee and Ahn, 2012; Stevens et al., 2020). However, researchers have not widely studied whether visual design features, augmented with AI prompt engineering, influence overt attention when considering other predictors of attention. Thus, we pose the following research question:

*RQ1: Will visual message design influence gaze duration and fixation level?*

The second research question focused on advertisement placement. Placement is a key factor, especially for outdoor advertising, but also on websites and other spatial advertising contexts. However, in the outdoor context, placement is difficult to manipulate experimentally. Some eye-tracking work has examined placement factors in field studies (Costa et al., 2019; Peker et al., 2021; Wilson and Casper, 2016), but the specific ad and the place where it is shown are typically confounded. One of the benefits of VR is that because the environment can be manipulated, we can randomly assign ads to places, thereby turning a previously quasi-experimental variable into an experimental one. With this in mind, we asked:

*RQ2: Will advertisement placement influence gaze duration and fixation count?*

## 4. Methods

The code and data are provided in a reproducibility package at [anonymized for review].

### 4.1. Participants

A total of 45 participants were recruited and received course credit for compensation. They were randomly assigned to either the sales sign counting or the free viewing conditions. Data from participants who didn’t properly understand the instructions of the study (*n* = 2) or had a below-normal vision (*n* = 2) were replaced, resulting in a final sample of 41 participants (*m*_*age*_ = 22.91; *sd*_*age*_ = 4.78), with 53% identifying as female). Approval for the study was obtained from the local Institutional Review Board, and all participants provided written informed consent before participation.

### 4.2. Materials and Equipment

#### 4.2.1. Advertisement Design Using Generative Artificial Intelligence (AI)

To design the advertisements, we first devised a list of 20 distinct commercial and 20 health topics through internet searches and from previous related work (Cho et al., 2024; Schmälzle et al., 2023). Then, we developed 2 versions of each advertisement - one intended to be more visually attention-grabbing than the other - using generative AI to augment our work (see Figure 3A). Specifically, each advertisement included a short text and an accompanying image. The text of the advertisements was either adopted from our previous studies (Cho et al., 2024; Schmälzle et al., 2023) or created with the help of ChatGPT via predefined prompts (e.g., Generate messages aimed at promoting handwashing; chatgpt.com). Then we used the Ideogram software service (ideogram.ai) to generate less attention-grabbing images, starting with predefined prompts (e.g., a boring and non attention-grabbing image that promotes sleep) and using the magic prompt feature to eliminate unrealistic features (e.g., disfigured hand, table missing a leg). For more attention-grabbing images, we used MidJourney ^3^ to generate candidate images through predefined prompts (e.g., imagine/An attention-grabbing health campaign that promotes sleep; www.midjourney.com).

**Figure 3:**
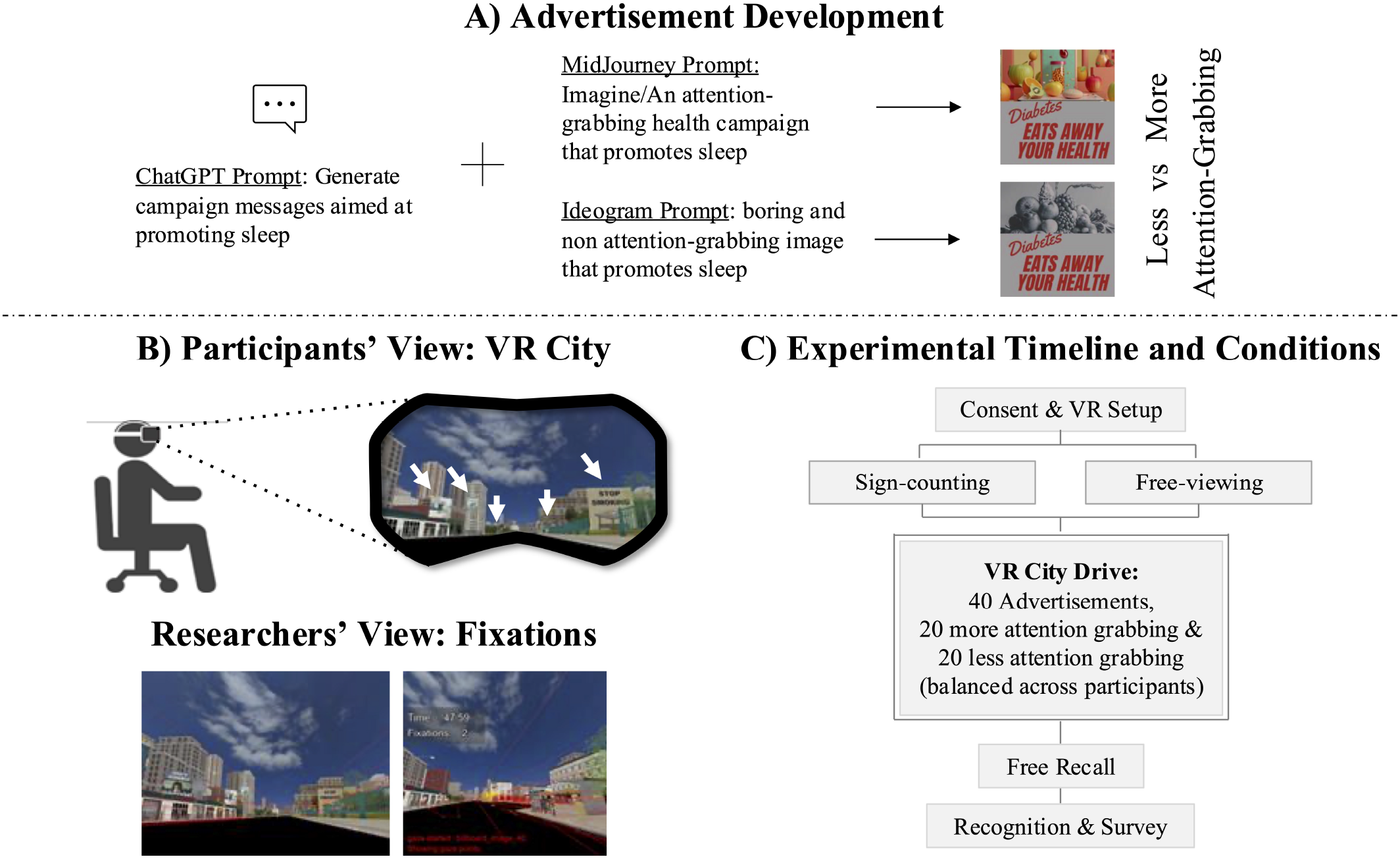
Advertisement Design and Experimental Procedures. The top (A) illustrates the predefined prompts we used to generate the text (via ChatGPT) and images (MidJourney for more attention-grabbing and Ideogram for less attention-grabbing) for the advertisements. The bottom left (B) demonstrates advertisement placement locations in the VR city (in white arrows) and the fixation data from the Vizard platform. Finally, the bottom right (C) outlines the experimental procedure.

Finally, on an 800 & 800 Canva billboard template, we placed the text and the less attention-grabbing image in distinct areas to enhance the readability of the text and selected the background color to match the image, minimizing color contrast. For the more attention-grabbing version of the advertisements, we replaced the ideogram-generated image with the Midjourney-generated image and kept everything else consistent. We selected Midjourney images that enhanced color contrast with the background. In total, we created 80 advertisements (40 more attention-grabbing and 40 less attention-grabbing). Our use of predefined AI prompts and consistently formatted advertisements on Canva ensured experimental control and standardization, while still reflecting realistic everyday variations.

#### 4.2.2. Virtual City and VR Head Mounted Display (HMD) with Eye-Tracking

Vizard’s Sightlab VR Pro Software (Worldviz.com) was used to run the experiment. We built the virtual city by importing the Classical City 3D model from Unity Asset (Classic City: Mobile, 2021) into Vizard. The environment included buildings, parks, and other objects typically found in a city. We modified the city to decrease the number of distractions and designated 40 advertisement placement positions on buildings and in parks throughout the city street (see Figure 3B). The placement locations were distributed relatively evenly to the left and right of the street, though the height, distance from the road, and angle of the advertisement varied slightly to enhance realism.

We randomized the order of the 40 advertisements for pairs of participants, and the pairs were exposed to the opposite visual design versions (i.e., more vs. less attention-grabbing) of the advertisements in the same order (e.g., participant 1: position index 1 - moving business more attention-grabbing, position index 2 - law firm less attention-grabbing, position index 3 - furniture store more attention-grabbing; participant 2: 1st Ad - moving business less attention-grabbing, 2nd Ad - law firm more attention-grabbing, 3rd Ad - furniture store less attention-grabbing). This method ensured that all advertisement versions were shown equally across the task manipulation.

We used the HP Reverb G2 Omnicept with eye-tracking capabilities for the VR head-mounted display. Participants drove straight down one city street using the right VR controller to decrease confusion and additional environmental distractions.

### 4.3. Experimental Procedures and Conditions

Participants came to the lab in person and reviewed the consent form. Once the participants signed the form, they completed a brief vision test and put on the VR headset. We calibrated the eye tracker in the VR headset, gave them the right VR controller, and explained that they will drive through a newly developed city, currently closed to traffic for maintenance. Those in the sale sign counting condition (*n* = 20) were asked to count the signs with the word “SALE,” while those in the free driving condition (*n* = 21) were asked to look around freely and relax. Then, we started the study, which lasted about 5 minutes.

After the participants had completed the study, we conducted a short interview about the virtual drive and asked them to list all the advertisements they remembered (free recall task). Then, the participants completed a post-study Qualtrics survey, which included items about demographics, perceived presence, virtual sickness, recognition test, and visual design manipulation check. Finally, we debriefed the participants.

### 4.4. Measures and Data Analyses

#### 4.4.1. Fixation and Gaze Duration

Through Python code on the Vizard platform, we collected fixation and total gaze duration information for each advertisement in real-time. We set the fixation threshold to 0.25s. This meant that when a participant’s eyes first landed on the advertisement, the code recorded the time as “gaze started: [advertisement name].” If the gaze was prolonged for 0.25 seconds, the code logged the time as “gaze fixated: [advertisement name].” The time the participant looked away from the advertisement was subsequently logged as “gaze ended: [advertisement name].” Then, we calculated the fixation count and gaze duration for each participant. Specifically, we calculated how many distinct times a participant fixated on an advertisement. Then, for an advertisement with at least one fixation, we calculated the total number of seconds each participant looked at the advertisement (gaze duration = “gaze ended” - “gaze started”).

#### 4.4.2. Recall and Recognition

During the post-study interview, participants were asked to share everything they remember seeing. We logged their response as 1 for recall if the participants remembered the advertisement’s message (e.g., I remember seeing an ad about HIV). If the participants only remembered an aspect of the advertisement (e.g., I remember seeing a picture of a bear) without any indication that they understood the advertisement’s purpose, then we coded it as 0 for recall. For the recognition test, we presented 44 sets of advertisements (2 versions for each advertisement and 4 additional distractors) and asked participants to select the advertisements they recognized via Qualtrics.

#### 4.4.3. Other Survey Measures

In addition to the main variables, we included items about perceived presence, virtual sickness, and visual design manipulation check. The spatial presence scale (Hartmann et al., 2015) asked participants to rate 5 items (e.g., *“I felt like I was transported to a different place.”*) from a scale of 1 (strongly disagree) to 5 (strongly agree). The virtual sickness measure (Kim et al., 2018) examined how much participants experienced 9 symptoms, including *“General Discomfort”* and *“Fatigue*,*”* from a scale of 1 (none) to 4 (severe). Finally, the visual design manipulation check asked participants to select the more attention-grabbing version of each advertisement. The result of the manipulation check is shown in Appendix Table A1.

#### 4.4.4. Data Analysis

We conducted the main data analysis in R. To test the effect of viewing behavior on memory (H1-H3), we fitted a mixed-effects logistic regression model for recall and recognition, with assigned task condition (sale sign counting vs. free driving), fixation level ^4^ (no fixation vs. fixation =1 vs. fixation >1), and total gaze duration (in seconds) as the main effects. The intercept was allowed to vary by advertisement to control for the baseline variance in the memorability of each advertisement. In addition, to examine how viewing behavior is influenced by task condition (H4), advertisement visual design (RQ1), and advertisement position (RQ2), we fit two mixed effects regression models: multinomial regression model for fixation level (mclogit R package; Elff (2022) and zero-inflated gamma regression for total gaze duration (glmmTMB R package; Brooks et al. (2017)). Task condition, advertisement placement (40 positions), and visual message design (less vs. more attention-grabbing) were the main effects, and the intercepts were allowed to vary by advertisement and participant to control for the baseline variance in the advertisements’ features and participants’ gaze behavior. Finally, we conducted an exploratory analysis of overt attention by advertisement placement (see Appendix B).

## 5. Results

### 5.1. Descriptive Analyses

Overall, participants reported a high level of spatial presence in the VR environment (*M*_*spatialpresence*_ = 3.73, *SD*_*spatialpresence*_ = 1.01). Additionally, participants reported minimal symptoms such as discomfort, fatigue, or eyestrain (*M*_*VRsymptoms*_ = 1.30, *SD*_*VRsymptoms*_ = .58). This indicated that the participants felt as if they were physically present in the city, and the technological aspects did not interfere much with their experience.

In addition, on average, participants fixated on about 70% (28/40) of the billboards (see Table 1). The average recall rate was around 8%, and the average recognition rate was 40%. These results suggested that the city environment included many city-typical elements (e.g., tall buildings and restaurant signs) that distracted people’s attention away from the advertisements.

**Table 1:** Average # of Ads. Fixated, Recalled, and Recognized per Person (Out of 40)

|  | Free Viewing | Sales Sign Counting | All Participants |
| --- | --- | --- | --- |
| Ads. Fixated | 65% (26/40) | 73% (29/40) | **70%** (28/40) |
| Recalled | 10% (4/40) | 8% (3/40) | **8%** (3/40) |
| Recognized | 45% (18/40) | 35% (14/40) | **40%** (16/40) |

### 5.2. Effect of Overt Attention and Task Condition on Memory (H1-H3)

Next, our regression models supported our hypotheses about the link between attention and memory (see Figure 4 and Table 2). As predicted, fixation level significantly predicted recall (*χ*^2^*(2)* = 15.78, *p* < .001) and recognition (*χ*^2^*(2)*= 23.28, *p* < .001), supporting H1a and H1b. In other words, fixating on an ad at least once (vs. 0 times) enhanced the likelihood of recall and recognition of the ad. Gaze duration also predicted recall (*χ*^2^*(1)* = 18.82, *p* < .001) and recognition (*χ*^2^*(1)* = 30.99, *p* < .001), supporting H2a and H2b. This result showed that the longer people looked at an advertisement, the more likely they would remember it.

**Table 2:** Effect of Overt Attention on Memory (Mixed Effects Logistic Regression Model)

|  | Estimate | S.E. | z | p-value |
| --- | --- | --- | --- | --- |
| **Dependent Variable: Recall** |  |  |  |  |
| Intercept | -4.54 | .38 | -12.00 | <.001 |
| Fixation Count: Fixation = 1 | 1.44 | .37 | 3.87 | <.001 |
| Fixation Count: Fixation > 1 | 1.12 | .41 | 2.72 | .0065 |
| Gaze Duration | .40 | .092 | 4.34 | <.001 |
| Task Condition: Free Viewing | .56 | .19 | 2.85 | .0044 |
| **Dependent Variable: Recognition** |  |  |  |  |
| Intercept | -1.54 | .15 | -10.50 | <.001 |
| Fixation Count: Fixation = 1 | .78 | .16 | 4.79 | <.001 |
| Fixation Count: Fixation > 1 | .58 | .20 | 2.88 | .0039 |
| Gaze Duration | .41 | .074 | 5.57 | <.001 |
| Task Condition: Free Viewing | .39 | .11 | 3.53 | <.001 |

Note. S.E. = Standard Error; Reference group for Fixation Count: "Fixation = 0"; Reference group for Task Condition: "Sale Sign Counting"; $R^2_{\text{Conditional Recall}} = .32$ ; $R^2_{\text{Marginal Recall}} = .19$ ; $R^2_{\text{Conditional Recognition}} = .18$ ; $R^2_{\text{Marginal Recognition}} = .13$

**Figure 4:**
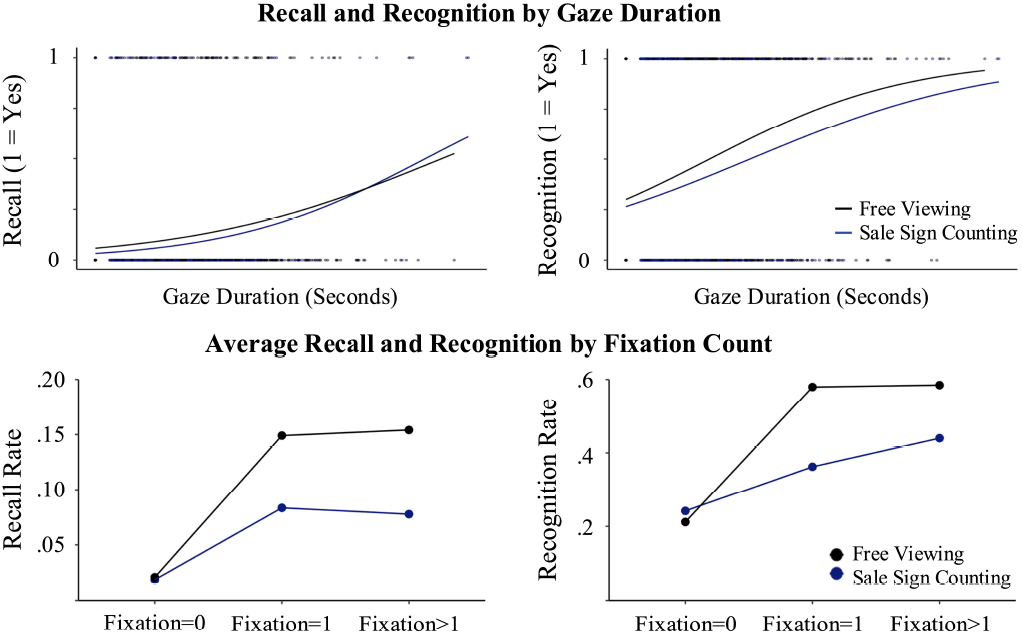
Effect of Overt Attention (Gaze Duration and Fixation Count) on Memory.

In addition, the task condition significantly influenced recall (*χ*^2^*(1)* = 8.12, *p* = .0044) and recognition (*χ*^2^*(1)* = 12.44, *p* < .001): Those assigned to the free driving condition were more likely to recall and recognize the advertisement than those in the sales sign counting condition see Figure 4 and Table 2). Thus, H3a and H3b were supported, suggesting that the parallel task interfered with information retention.

### 5.3. Effects of Task Condition, Ad Placement, and Message Design on Attention (H4, RQ1-2)

Furthermore, the results shed interesting insights into the predictors of overt attention (see Figure 5 and Tables 3-4 in the Appendix). Interestingly, those in the sale sign counting condition were more likely to fixate on an advertisement once (vs. 0 times) compared to their counterparts in the free driving condition (*estimate* = −.67, *z* = −2.28, *p* = .023). Task condition did not influence whether participants fixated on the advertisement multiple times (vs. 0 times; *estimate* = −.28, *z* = −.82, *p* = .41). Thus, H4a was not supported. However, those in the free driving condition looked at advertisements longer than those in the sale sign counting condition (*estimate* = .41, *z* = 6.04, *p* < .001), supporting H4b. One likely explanation for this discrepancy between fixation and gaze duration is that participants in the sale sign counting condition looked at advertisements long enough to see if it was a “SALE” sign, while those in the free viewing condition freely engaged with the advertisements.

**Figure 5:**
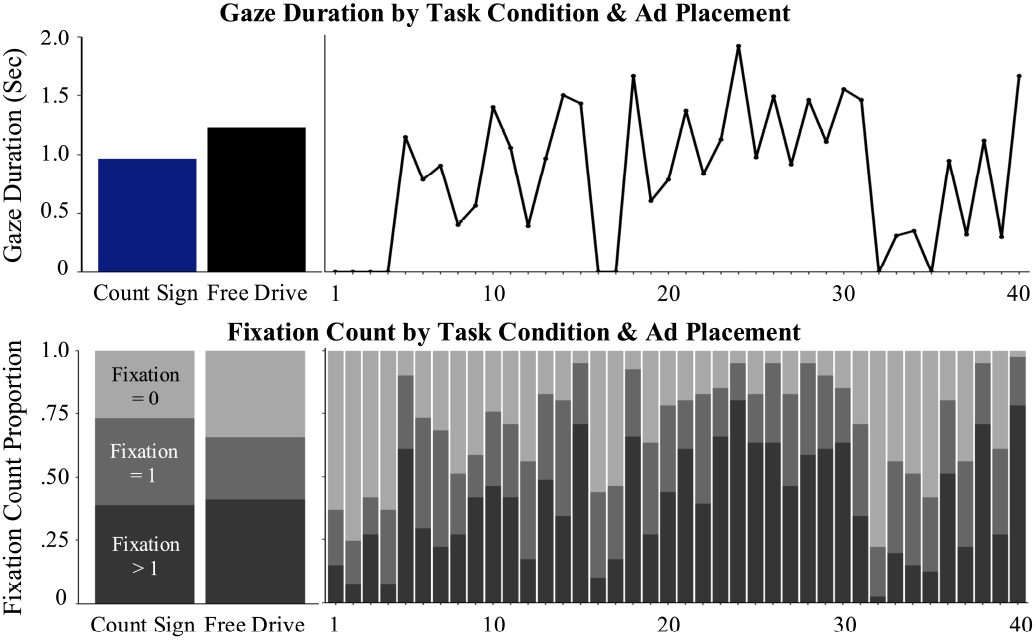
Gaze Duration (Median) and Fixation Count by Task Condition and Ad Placement.

For other predictors, advertisement placement significantly predicted fixation level (see Table 3 in the appendix) and gaze duration (*χ*^2^*(39)* = 166.46, *p* < .001). In other words, certain locations in the virtual environment attracted more attention than others. Visual message design did not significantly influence fixation or gaze duration when task condition and advertisement placement were held constant.

## 6. Discussion

This study tested four hypotheses and two research questions about the causal chain of advertising effects, starting from ads in the visual environment, attention to/reception of those messages, and finally memory for the messages. Our main novel contributions are that we combined the advantages of eye-tracking with immersive VR to rigorously examine the factors that influence attention and retention and used generative AI for advertisement design. Below, we discuss the key findings as well as the broader theoretical and practical implications of this approach for advertising research.

Our findings supported the main hypotheses regarding the effects of overt attention and attentional tasks on memory performance. Increased fixation and total gaze duration explain both recall and recognition rates, confirming H1 and H2. These results reinforce the established link between attention and memory as measured by eye-tracking and memory tests (Bott et al., 2017; Chang and Choi, 2014; Schmälzle et al., 2023). Moreover, participants in the free driving condition demonstrated superior recall and recognition compared to those engaged in the sales sign counting task, supporting H3. This finding aligns with previous research on the detrimental effects of distraction on retention (Fernandes and Moscovitch, 2000; Jacoby et al., 1989).

Analysis of factors influencing overt attention also yielded noteworthy results. As predicted by H4b, participants in the free driving condition exhibited longer gaze durations on advertisements compared to those in the sales sign counting condition, consistent with previous research on attention-demanding tasks and gaze behavior (Bang and Wojdynski, 2016; Wang et al., 2012). Contrary to H4a, however, participants in the sales sign counting condition were more likely to fixate at least once on non-target advertisements. This discrepancy between fixation and gaze duration measures likely reflects different attentional processes: brief fixations on non-target ads may result from active searching for “SALE” signs, while longer gazes in the free driving condition suggest intrinsic interest in the billboards (Wolfe and Horowitz, 2017).

Regarding the influence of ad placement on attention, the positioning of billboards within the city significantly influenced both fixation and total gaze duration measures. This aligns with related studies on ad placement, from website location to real-world billboard positioning (AlKheder, 2024; Resnick and Albert, 2014; Wilson and Casper, 2016). Research on billboard placement has confirmed that factors such as strategic location (e.g., Times Square in NYC or Piccadilly Circus in London), traffic volume (e.g., busy intersections), audience characteristics, and numerous other variables (e.g., proximity to hospitals, schools) influence the effectiveness and ROI of outdoor advertising (Bhargava et al., 1993; Donthu, 1995; Rossiter and Bellman, 2005; Wilson and Till, 2008). Through experimental control with randomized ad assignment to positions, we isolated the effect of placement more precisely. This allowed us to quantify the value of specific locations based on their ability to attract audience attention, effectively assigning an attentional real estate value to every corner of our simulated city (see Appendix B).

Lastly, visual design characteristics (AI-generated, less vs. more attention-grabbing) did not influence gaze when all other predictors were kept constant. However, the post-experimental data in which participants were asked (via a forced choice) to select the more attention-grabbing ad from the two versions supported our manipulation (see Appendix A). We present a few potential explanations for these findings. First, research has found discrepancies between self-reported attention and objective eye-tracking measures (Beuckels et al., 2021). The discrepancy suggests that the two methodologies could measure slightly different things. The self-report could measure people’s evaluations of the messages after careful inspection of the two versions side-by-side, whereas eye-tracking measures the immediate biological response to a version of the stimuli without seeing the other version. Furthermore, the difference between the less and more attention-grabbing versions may not have been strong enough to impact real-time overt attention, especially when presented with other predicting variables.

### 6.1. Theoretical and Methodological Contributions and Practical Implications

This section discusses the theoretical contributions and practical implications together, as theoretical advances in causal mechanism identification directly translate into practical applications in advertising.

#### 6.1.1. Self-Directed Attention and Incidental Memory

Our study significantly advances the theoretical understanding of overt visual attention and incidental memory by elucidating how parallel tasks and environmental distractions impact advertising effectiveness. In real-world settings like cities, a myriad of distractions can affect one’s attention and memory processes. Our findings show that parallel tasks significantly decreased prolonged attention and overall retention of an ad. This highlights the cognitive interference caused by the parallel tasks, providing a more nuanced understanding of our divided attention in highly realistic everyday contexts. By examining the causal chain of advertising effects, we explored the sequence from the presence of ads in the visual environment, exposure through the attention to and reception of these messages, and ultimately to memory retention. This comprehensive approach offers in-depth insights into the cognitive mechanisms underlying the reception and retention of advertising messages, highlighting the critical stages where environmental factors can disrupt attention and memory processes.

Additionally, our results indicate that traditional ad features (e.g., message designs) were less impactful in a distraction-prone environment like a city. While generative AI can produce high-quality images and messages, these may not be sufficient in noisy settings. Thus, further theoretical exploration is required to understand what makes messages “attention grabbing” in such contexts. Practically, advertisers must be mindful of these findings, acknowledging the limitations of the ad features in environments full of distractions and focusing on strategies that retain attention amidst distractions. For example, practitioners could combine generative AI’s capability to create visually appealing messages with physical locations of the advertisements to optimize people’s initial attention and their future retention of the advertisement.

#### 6.1.2. Implications of Method-Theory Synergy

An important contribution of our approach is the integration of biometric research, experimental rigor, and realism afforded by the VR city paradigm. Traditionally, theoretical, mechanism-focused research on advertising effects has faced methodological barriers like the tradeoff between experimental control and realism (De Pelsmacker, 2021), and the inherent limitations of biometric research in lab settings (Potter and Bolls, 2012). VR technology overcomes these barriers, enabling even studies in yet-to-be-built cities or fantasy environments. Additionally, the VR-based approach has advantages over field studies by allowing experimental manipulation of billboards and environmental control. This capability is essential for mechanism identification, paving the way for “causal inference in generalizable environments” (Miller et al., 2019). In this sense, what may initially appear as a methodological advancement has important theoretical consequences as it brings us closer to understanding the causal factors that drive advertising success (Greenwald, 2012).

For instance, many biometric advertising research paradigms force exposure by placing people in front of screens and asking them to view each message. These research paradigms allow studying reception processes, but they fail to represent incidental exposure, which allows people to self-select messages to pay attention to based on a complex interaction between environmental, user-sided, and message-related variables. However, real-world advertising hinges on incidental exposure and reception. Thus, with VR, we can experimentally control theorized variables while capturing biometric variables in situ. This capability opens up a whole new avenue of research at the intersection of real-life exposure contexts and VR-mediated biometric research.

In addition, measuring the effect of outdoor advertisements on the audience has long been a critical focus (Bloom, 2000), and our results show promising advancements in this area. While current methods, such as smartphone GPS data, can provide a broad overview of potential audiences, they lack granularity, like observing traffic from a helicopter. Our approach allows us to determine whether a specific driver actually views a billboard they pass, providing deeper insights into the causal mechanisms of advertising effectiveness. And compared to inventive experimental research that manipulated outdoor ads (King and Tinkham, 1989), the current approach is far more cost-effective.

Further synergy between theory and method, and between advertising theory and practice, arises from another important consideration. In the study presented here, VR served as a kind of experimental petri-dish, or a realistic simulation of the world that included key aspects like spatial vision and free exploration. However, as the internet advertising market evolves or expands towards the Metaverse (which seems likely given the investments of key players like Meta, Apple, and Tencent), VR-based environments will become the new advertising environments in their own right (Kim, 2021). Our theoretical arguments about exposure to messages, attention/reception, and subsequent memory also extend this context.

Perhaps most relevant for advertising practice: VR-based paradigms with integrated eye trackers have predictive potential. Eye-tracking data streams are comparatively easy to collect, featurize, and integrate into predictive pipelines (Coronel et al., 2021). Comparable predictive or machine-learning approaches have revolutionized the digital advertising ecosystems. Given the rapid advancement of VR technologies and the metaverse, we can expect a similar development for VR-based content and advertising revenue models. This development enables practices such as next-gaze prediction, attention-based and dynamic ad pricing, or repeated ad delivery to reach the optimal level between ad exposure and the target audience’s attention.

These considerations suggest that the type of VR-based research presented here may soon evolve into a core addition to the advertising researchers’ toolbox. So far, we have seen the field embrace tools like focus groups, survey panels, or online experimentation. Although the use of VR-based simulacra for pretesting is not yet fully formalized, it is very much possible - and in some cases (e.g., real estate marketing), already a reality.

#### 6.1.3. Ethical Considerations

Of course, any research paradigm involving biometric measurements raises many ethical questions - ranging from user privacy, ownership of biometric data, and the broader issue of protection from manipulation. As researchers, we can benefit from the deeper insights biometric methods afford, which can also help to better understand and improve the effectiveness of social marketing and health communication efforts. However, given that the risks and benefits of new technologies correlate, a broader debate about biometrics-based (particularly eyetracking) advertising seems indicated (Farahany, 2023; Ravi, 2017).

### 6.2. Future Research Directions

Here, we discuss promising avenues for future research. First, researchers could modify the features of the city paradigm to further enhance ecological validity and study other theoretical concepts. We deliberately had participants drive down a straight street rather than navigating the entire city environment to reduce potential confounding variables such as idiosyncratic navigation behavior. Future studies can ask participants to navigate the environment and capture eye-tracking metrics along the way. Through this modification, researchers can examine the role of spatial navigation and place conditioning, which has long been known to be intimately interwoven with memory systems in the brain (Eichenbaum, 2017). Along a similar line of reasoning, researchers can add additional city-typical elements such as dynamic traffic and people or other realistic motivations or goals for the participants (e.g., needing to buy something) to enhance ecological validity even more and study other interesting theoretical variables.

Next, our paradigm can be expanded to contexts beyond a city setting. Existing studies have already examined contexts such as highways or shopping malls. Other promising environments for advertising research include airports, metro/railway stations, or larger to-be-built architectural projects like new theaters and other public venues. Our work on quantifying attentional real estate value based on ad locations can especially provide valuable insights through these simulated settings.

Furthermore, future research can examine theoretical variables beyond attention and memory. For instance, it would be promising to incorporate actual behavior, such as purchase decisions in shopping contexts. Doing so would allow the examination of the effect of advertising factors on behavioral outcomes and also provide a real-world anchoring to the virtual reality experience. Motivational variables, such as hunger or other need states, could also provide interesting insights into advertising outcomes (Stockburger et al., 2009).

Lastly, the flexibility of VR opens up creative avenues for studying ad delivery and execution. Digital billboards with dynamic content and attention-grabbing features can be easily simulated (e.g., light pulses as eye-catchers; Melendrez-Ruiz et al. (2021)). Future developments could also incorporate embodied AI agents as artificial influencers, directing attention to ads or engaging users in real-time ad-related conversations (Lim et al., 2024).

## 7. Summary and Conclusions

This study combined the use of VR and eye-tracking to study attention to and retention of advertisements. The use of VR enabled the creation of a key advertising context - an urban city environment - and the use of eye-tracking enabled us to objectively assess whether people actually looked at the billboards they passed. Our results show that this information about actual exposure, which comprises overt attention to facilitate message intake, robustly explains message memory. Moreover, we demonstrate the strong influence of ad placement. This approach opens up exciting possibilities to study theoretical mechanisms of advertising, practically test ad copy in realistic contexts, and fruitfully combine biometric markers like fixations and viewing time with traditional advertising metrics like reach, frequency, and impressions.

## Data Availability Statement

This study’s data will be available on GitHub once the manuscript is accepted.

## Footnotes

3 We used two different AI-image generation tools because MidJourney exhibited bias toward visually appealing images. No matter what prompts we used, MidJourney’s “less attention-grabbing” images were still very visually appealing.

4 As expected, fixation count and total gaze duration were highly correlated (*r* = .80). To decrease multicollinearity, we used fixation count categories rather than the raw counts. We split fixation count into 3 categories based on the median fixation count (fixation = 1).

