## Appendix for "Using VR and eye-tracking to study attention to and retention of AI-generated ads in outdoor advertising environments"

### Tables 3 and 4

**Table 3. Predictors of Gaze Duration (Mixed Effects Gamma Regression Model)**

| | ***Estimate*** | ***S.E.*** | ***z*** | ***p-value*** |
|---|---|---|---|---|
| Intercept | -.079 | .16 | *-.50* | .62 |
| Task Condition: Free Viewing | .41 | .068 | *6.04* | **<.001** |
| Visual Ad. Design: More Attention-Grabbing | .021 | .034 | *.62* | .54 |
| Ad. Placement: Position 2 | .20 | .24 | *.84* | .40 |
| Ad. Placement: Position 3 | .21 | .21 | *1.03* | .30 |
| Ad. Placement: Position 4 | -.056 | .21 | *-.26* | .79 |
| Ad. Placement: Position 5 | .39 | .18 | *2.21* | **.027** |
| Ad. Placement: Position 6 | .51 | .18 | *2.76* | **.0058** |
| Ad. Placement: Position 7 | .45 | .19 | *2.44* | **.015** |
| Ad. Placement: Position 8 | .35 | .19 | *1.78* | .075 |
| Ad. Placement: Position 9 | .47 | .19 | *2.46* | **.014** |
| Ad. Placement: Position 10 | .43 | .18 | *2.38* | **.017** |
| Ad. Placement: Position 11 | .44 | .18 | *2.36* | **.018** |
| Ad. Placement: Position 12 | -.018 | .19 | *-.093* | .93 |
| Ad. Placement: Position 13 | .10 | .18 | *.57* | .57 |
| Ad. Placement: Position 14 | .23 | .18 | *1.29* | .20 |
| Ad. Placement: Position 15 | .30 | .18 | *1.71* | .088 |
| Ad. Placement: Position 16 | -.047 | .20 | *-.23* | .82 |
| Ad. Placement: Position 17 | .13 | .20 | *.63* | .53 |
| Ad. Placement: Position 18 | .44 | .18 | *2.51* | **.012** |
| Ad. Placement: Position 19 | -.034 | .19 | *-.18* | .86 |
| Ad. Placement: Position 20 | .012 | .18 | *.065* | .95 |
| Ad. Placement: Position 21 | .30 | .18 | *1.65* | .10 |
| Ad. Placement: Position 22 | .18 | .18 | *1.00* | .32 |
| Ad. Placement: Position 23 | .22 | .18 | *1.25* | .21 |
| Ad. Placement: Position 24 | .63 | .18 | *2.56* | **<.001** |
| Ad. Placement: Position 25 | .42 | .18 | *2.31* | **.021** |
| Ad. Placement: Position 26 | .50 | .18 | *2.82* | **.0049** |
| Ad. Placement: Position 27 | .088 | .18 | *.49* | .63 |
| Ad. Placement: Position 28 | .29 | .18 | *1.62* | .11 |
| Ad. Placement: Position 29 | .34 | .18 | *1.92* | .054 |
| Ad. Placement: Position 30 | .61 | .18 | *3.39* | **<.001** |
| Ad. Placement: Position 31 | .43 | .18 | *2.37* | **.018** |
| Ad. Placement: Position 32 | -.14 | .25 | *-.55* | .58 |
| Ad. Placement: Position 33 | -.079 | .19 | *-.41* | .68 |
| Ad. Placement: Position 34 | .022 | .20 | *.11* | .91 |

| | | | | |
|---|---|---|---|---|
| Ad. Placement: Position 35 | -.18 | .20 | *-.89* | .37 |
| Ad. Placement: Position 36 | .16 | .18 | *.87* | .39 |
| Ad. Placement: Position 37 | -21 | .19 | *-1.07* | .28 |
| Ad. Placement: Position 38 | .12 | .18 | *.69* | .49 |
| Ad. Placement: Position 39 | -.27 | .19 | *-1.42* | .15 |
| Ad. Placement: Position 40 | .45 | .18 | *2.54* | **.011** |

*Note. S.E. = Standard Error; Estimate in log-odds scale; Reference group for Task Condition: "Sale Sign Counting"; Reference group for Ad. Placement: "Position 1"; Reference group for Visual Ad. Design: "Less Attention-Grabbing"; $R^2$ Conditional* = .34; *$R^2$ Marginal* = .21

**Table 4. Predictors of Fixation Count (Mixed Effects Multinomial Regression Model)**

| | Fixation = 1 vs. 0 | | | | Fixation > 1 vs. 0 | | | |
|---|---|---|---|---|---|---|---|---|
| | *Est.* | *S.E.* | *z* | *p-value* | *Est.* | *S.E.* | *z* | *p-value* |
| Intercept | -.80 | .46 | *-1.73* | .084 | -1.55 | .53 | *-2.90* | **.0037** |
| Task Condition: Free Viewing | -.67 | .30 | *-2.28* | **.023** | -.28 | .34 | *-.82* | .41 |
| Visual Ad. Design: More Attention-Grabbing | .014 | .14 | *.10* | .92 | .16 | .15 | *1.08* | .28 |
| Ad. Placement: Position 2 | -56 | .61 | *-92* | .36 | -.88 | .78 | *-1.13* | .26 |
| Ad. Placement: Position 3 | -.34 | .63 | *-53* | .59 | .68 | .62 | *1.10* | .27 |
| Ad. Placement: Position 4 | .30 | .56 | *.54* | .59 | -.70 | .79 | *-.89* | .37 |
| Ad. Placement: Position 5 | 2.49 | .73 | *3.43* | **<.001** | 3.70 | .74 | *5.00* | **<.001** |
| Ad. Placement: Position 6 | 1.75 | .58 | *3.01* | **.0026** | 1.81 | .65 | *2.79* | **.0053** |
| Ad. Placement: Position 7 | 1.65 | .56 | *2.94* | **.0032** | 1.31 | .66 | *1.99* | **.046** |
| Ad. Placement: Position 8 | .44 | .58 | *.76* | .45 | 1.02 | .62 | *1.63* | .102 |
| Ad. Placement: Position 9 | .26 | .62 | *.41* | .68 | 1.65 | .60 | *2.74* | **.0062** |
| Ad. Placement: Position 10 | 1.46 | .61 | *2.40* | **.016** | 2.37 | .63 | *3.75* | **<.001** |
| Ad. Placement: Position 11 | 1.30 | .59 | *2.19* | **.029** | 2.08 | .63 | *3.33* | **<.001** |
| Ad. Placement: Position 12 | 1.05 | .55 | *1.91* | .056 | .62 | .67 | *.93* | .35 |
| Ad. Placement: Position 13 | 2.01 | .64 | *3.16* | **.0016** | 2.85 | .67 | *4.27* | **<.001** |
| Ad. Placement: Position 14 | 2.15 | .61 | *3.56* | **<.001** | 2.24 | .67 | *3.36* | **<.001** |
| Ad. Placement: Position 15 | 3.05 | .89 | *3.41* | **<.001** | 4.57 | .89 | *5.12* | **<.001** |
| Ad. Placement: Position 16 | .63 | .55 | *1.15* | .25 | -.24 | .74 | *-.33* | .75 |
| Ad. Placement: Position 17 | .46 | .56 | *.82* | .41 | .39 | .66 | *.59* | .56 |
| Ad. Placement: Position 18 | 2.69 | .79 | *3.42* | **<.001** | 4.08 | .79 | *5.16* | **<.001** |
| Ad. Placement: Position 19 | 1.15 | .57 | *2.02* | **.044** | 1.30 | .64 | *2.04* | **.041** |
| Ad. Placement: Position 20 | 1.71 | .61 | *2.82* | **.0048** | 2.46 | .64 | *3.82* | **<.001** |
| Ad. Placement: Position 21 | 1.34 | .66 | *2.03* | **.043** | 2.93 | .64 | *4.55* | **<.001** |
| Ad. Placement: Position 22 | 2.23 | .62 | *3.58* | **<.001** | 2.64 | .67 | *3.91* | **<.001** |
| Ad. Placement: Position 23 | 1.63 | .70 | *2.34* | **.019** | 3.29 | .68 | *4.87* | **<.001** |
| Ad. Placement: Position 24 | 2.53 | .93 | *2.72* | **.0065** | 4.80 | .89 | *5.39* | **<.001** |
| Ad. Placement: Position 25 | 1.44 | .68 | *2.12* | **.034** | 3.15 | .66 | *4.78* | **<.001** |
| Ad. Placement: Position 26 | 3.25 | .88 | *3.69* | **<.001** | 4.50 | .90 | *5.03* | **<.001** |
| Ad. Placement: Position 27 | 2.10 | .63 | *3.32* | **<.001** | 2.84 | .67 | *4.26* | **<.001** |
| Ad. Placement: Position 28 | 3.41 | .88 | *3.88* | **<.001** | 4.38 | .90 | *4.88* | **<.001** |
| Ad. Placement: Position 29 | 2.44 | .73 | *3.35* | **<.001** | 3.60 | .74 | *4.86* | **<.001** |
| Ad. Placement: Position 30 | 1.70 | .69 | *2.47* | **.013** | 3.24 | .68 | *4.79* | **<.001** |
| Ad. Placement: Position 31 | 1.43 | .58 | *2.47* | **.014** | 1.85 | .63 | *2.92* | **.0036** |
| Ad. Placement: Position 32 | -.44 | .59 | *-.74* | .46 | -2.06 | 1.13 | *-1.82* | .069 |
| Ad. Placement: Position 33 | 1.03 | .56 | *1.85* | .064 | .85 | .65 | *1.30* | .19 |
| Ad. Placement: Position 34 | .87 | .55 | *1.57* | .12 | .47 | .68 | *.69* | .49 |

| | | | | | | | | |
|---|---|---|---|---|---|---|---|---|
| Ad. Placement: Position 35 | .42 | .56 | *.75* | .45 | -.072 | .70 | *-10* | .92 |
| Ad. Placement: Position 36 | 1.71 | .63 | *2.70* | **.0069** | 2.72 | .65 | *4.17* | **<.001** |
| Ad. Placement: Position 37 | .96 | .56 | *1.72* | .086 | .87 | .64 | *1.35* | .18 |
| Ad. Placement: Position 38 | 3.07 | .90 | *3.43* | **<.001** | 4.65 | .90 | *5.18* | **<.001** |
| Ad. Placement: Position 39 | 1.06 | .56 | *1.89* | .059 | 1.20 | .63 | *1.91* | .057 |
| Ad. Placement: Position 40 | 3.48 | 1.15 | *3.02* | **.0025** | 5.41 | 1.14 | *4.75* | **<.001** |

*Note. Est. = Estimate; S.E. = Standard Error; Estimate in log-odds scale; Reference group for Task Condition: "Sale Sign Counting"; Reference group for Ad. Placement: "Position 1"; Reference group for Visual Ad. Design: "Less Attention-Grabbing"; Nagelkerke's* $R^2 = .38$

### Appendix A: How Well Did the Visual Design Manipulation Work?

In addition to the main analyses, we asked participants to select the more attention-grabbing version of an advertisement (see Table A1). The majority of the participants (>50%) selected the correct more attention-grabbing version of most advertisements.

**Table A1. % of Participants who Correctly Selected the More Attention-Grabbing Ad.**

| Commercial Ad Topic | % of Participants | Health Ad Topic | % of Participants |
|---|---|---|---|
| Burger Restaurant | 100% | Healthy Diet Promotion | 100% |
| Travel Agency | 98% | Handwashing Promotion | 95% |
| Museum | 98% | Smoking Prevention | 93% |
| Law Firm | 95% | Jogging Promotion | 93% |
| Car Insurance | 95% | Diabetes Prevention | 90% |
| Pet Store | 95% | Sun Protection | 90% |
| Cosmetics | 93% | Employee Health Awareness | 88% |
| Grocery Store | 93% | Sleep Promotion | 88% |
| Charity Donation | 90% | Visit Doctor | 83% |
| Hotel | 88% | Vaccination Promotion | 78% |
| Pottery Class | 88% | Mental Health Promotion | 78% |
| Furniture Store | 85% | Putting on Seatbelt | 76% |
| Smartphone Brand | 83% | Binge Drinking Prevention | 73% |
| Moving Business | 83% | Distracted Driving Prevention | 73% |
| Carpet Cleaning Service | 78% | HIV Prevention | 73% |
| Clothes Sale | 76% | Decrease Caffeine Intake | 73% |
| Pizza Restaurant | 68% | Save the Great Lakes | 73% |
| Movie Premier | 59% | Save Polar Bears | 68% |
| Brunch Restaurant | 54% | Recycle Promotion | 61% |
| Coffee Shop | 39% | Drug Abuse Prevention | 51% |

#### Appendix B: Exploratory Analysis of Advertisement Placement

In addition to the main analysis, we conducted an exploratory analysis of the 40 advertisement placements in the virtual city (see Figure B1).

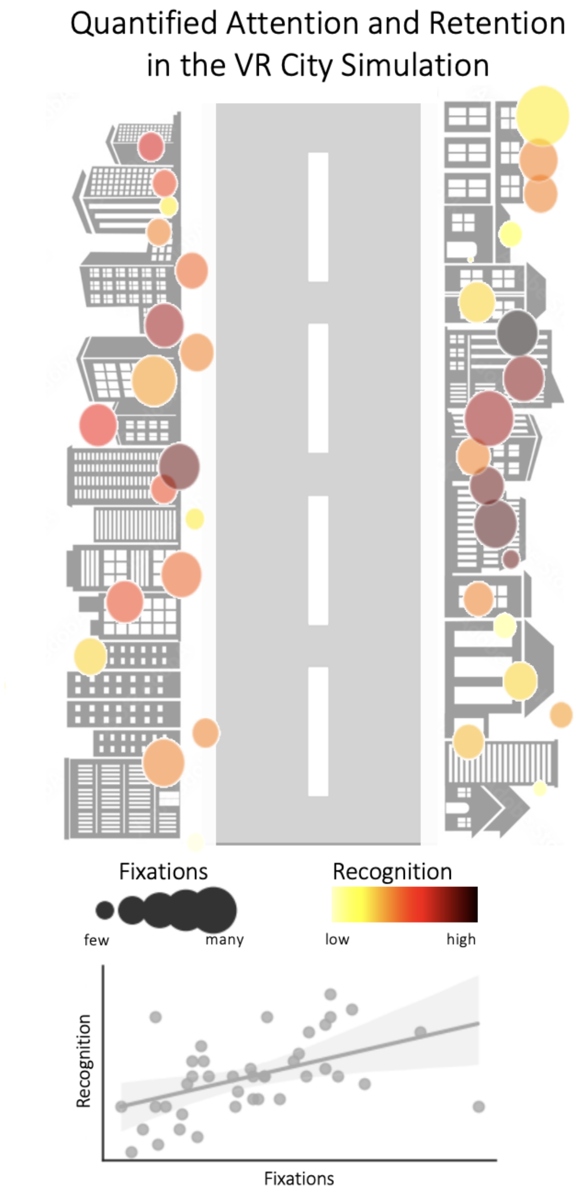


***Figure B1: Quantifying Attention and Retention in Their Spatio-Temporal Context along the Driving Route Through the Virtual City.*** *Bubbles are shown at the relative vertical/horizontal position along the city drive, with bubble size representing the total number of fixations per*

*location (standardized) and bubble color representing the recognition rate. The scatterplot in the lower panel shows that these aggregate metrics are positively correlated ($r = .48$, $p = .0016$).*
